# A cocktail containing two synergetic antibodies broadly neutralizes SARS-CoV-2 and its variants including Omicron BA.1 and BA.2

**DOI:** 10.1101/2022.04.26.489529

**Authors:** Xinghai Zhang, Feiyang Luo, huajun Zhang, Hangtian Guo, Junhui Zhou, Tingting Li, Shaohong Chen, Shuyi Song, Meiying Shen, Yan Wu, Yan Gao, Xiaojian Han, Yingming Wang, Chao Hu, Yuchi Lu, Wei Wang, Kai Wang, Ni Tang, Tengchuan Jin, Chengyong Yang, Guofeng Cheng, Haitao Yang, Aishun Jin, Xiaoyun Ji, Rui Gong, Sandra Chiu, Ailong Huang

## Abstract

Neutralizing antibodies (NAbs) can prevent and treat infections caused by severe acute respiratory syndrome coronavirus 2 (SARS-CoV-2). However, continuously emerging variants, such as Omicron, have significantly reduced the potency of most known NAbs. The selection of NAbs with broad neutralizing activities and the identification of conserved critical epitopes are still urgently needed. Here, we identified an extremely potent antibody (55A8) by single B-cell sorting from convalescent SARS-CoV-2-infected patients that recognized the receptor-binding domain (RBD) in the SARS-CoV-2 spike (S) protein. 55A8 could bind to wild-type SARS-CoV-2, Omicron BA.1 and Omicron BA.2 simultaneously with 58G6, a NAb previously identified by our group. Importantly, an antibody cocktail containing 55A8 and 58G6 (2-cocktail) showed synergetic neutralizing activity with a half-maximal inhibitory concentration (IC_50_) in the picomolar range *in vitro* and prophylactic efficacy in hamsters challenged with Omicron (BA.1) through intranasal delivery at an extraordinarily low dosage (25 μg of each antibody daily) at 3 days post-infection. Structural analysis by cryo-electron microscopy (cryo-EM) revealed that 55A8 is a Class III NAb that recognizes a highly conserved epitope. It could block angiotensin-converting enzyme 2 (ACE2) binding to the RBD in the S protein trimer via steric hindrance. The epitopes in the RBD recognized by 55A8 and 58G6 were found to be different and complementary, which could explain the synergetic mechanism of these two NAbs. Our findings not only provide a potential antibody cocktail for clinical use against infection with current SARS-CoV-2 strains and future variants but also identify critical epitope information for the development of better antiviral agents.

## Introduction

The coronavirus disease 2019 (COVID-19) pandemic caused by severe acute respiratory syndrome coronavirus 2 (SARS-CoV-2)^1^ has lasted for over two years. Although several antiviral agents (e.g., vaccines and neutralizing antibodies (NAbs)) have been authorized for the prevention and treatment of SARS-CoV-2 infection, the rapid emergence of variants of concern (VOCs), such as the Omicron variant, significantly reduces the efficacies of these antiviral agents and worsens the clinical outcomes achieved with them^2, 3^. Furthermore, it was recently reported that no authorized NAbs adequately cover all sublineages of the Omicron variant, except for the recently authorized agent LY-CoV1404 (bebtelovimab)^4^. Therefore, the identification of NAbs with broad neutralizing activities and conserved critical epitopes is still urgently needed.

SARS-CoV-2 belongs to the subgenus Sarbecovirus of the genus Betacoronavirus^5^. The SARS-CoV-2 spike (S) protein is a typical class I viral fusion protein that contains a surface subunit (S1) and a transmembrane subunit (S2). The entry of SARS-CoV-2 into host cells relies on the interaction between the receptor-binding domain (RBD) in S1 and its obligate receptor, angiotensin-converting enzyme 2 (ACE2)^6^. Structural determination of the RBD/ACE2 complex revealed that the SARS-CoV-2 RBD contains a receptor-binding motif (RBM) that is the core region for recognition^7^. The RBD in the S protein has two distinct conformations, with “up” or “open” representing the receptor-accessible state and “down” or “closed” representing the receptor-inaccessible state^8–10^. Because the RBD is critical for viral entry and is highly antigenic^11^, it is an attractive target for drug and vaccine development. The pattern of recognition of the RBD by NAbs determines the neutralizing breath and potency, which is important for the design and optimization of antibodies/or antibody cocktails as therapeutic agents and potential S protein-based immunogens for vaccines^12^. Currently, according to the epitopes defined by structural information for the complexes formed by NAbs and S proteins, antibodies can be classified into at least four categories (Classes I-IV): (1) NAbs (e.g., CB6) that recognize the RBM in only the up RBD conformation and block ACE2 by mimicking ACE2 binding; (2) ACE2-blocking NAbs that bind both the up and down conformations of the RBD and can contact adjacent RBDs (e.g., REGN10933); (3) NAbs that bind outside the ACE2 site of the RBD (non-RBM) and recognize both the up and down conformations of the RBD (e.g., S309 and LY-CoV1404); and (4) NAbs that bind far away from the ACE2 site by recognizing only the up conformation of the RBD and do not block ACE2 (e.g., CR3022 and ADG-2)^4, 12–14^. In general, the majority of mutations in VOCs occur in the RBM, which significantly reduces or completely eliminates the efficacies of Class I and II NAbs. In contrast, Class III and IV NAbs are more resistant to these mutations^14^. Notably, although S309 (also known as sotrovimab), a NAb that received an emergency use authorization (EUA), could still neutralize Omicron BA.1 and BA.1+R346K, its neutralizing activity against Omicron BA.2 was dramatically reduced. As mentioned above, only LY-CoV1404 can efficiently neutralize all sublineages of the Omicron variant.

In a previous study, we isolated a panel of NAbs with strong neutralizing activities against SARS-CoV-2^15^. Among these NAbs, 58G6 was recently found to inhibit all authentic VOCs, including the Delta and Omicron variants, with half-maximal inhibitory concentration (IC_50_) values of 1.69 ng/ml and 54.31 ng/mL, respectively. Importantly, this NAb has been shown to provide protection against the Delta and Omicron variants *in vivo* (under review). Here, we report a newly identified NAb termed 55A8 that can broadly neutralize wild-type (WT) SARS-CoV-2 (defined as SARS-CoV-2 unless otherwise indicated), as well as Omicron BA.1 and BA.2 with extreme potency. In addition, 55A8 and 58G6 showed synergetic effects both *in vitro* and *in vivo* that enhanced the SARS-CoV-2 neutralizing potency and breadth. Structural analysis revealed that 55A8 is a Class III NAb that recognizes a conserved epitope. Therefore, a cocktail of 55A8 and 58G6 (2-cocktail) could be developed to combat currently circulating VOCs (e.g., Omicron) and possible emerging VOCs in the future.

## Results

### 55A8 strongly binds to S proteins from SARS-CoV-2 and different variants

55A8 was isolated by sorting single B cells producing antibodies with potent neutralizing activity against SARS-CoV-2 and the Beta (B.1.135) authentic virus from convalescent patients previously infected with SARS-CoV-2^16^. Here, we showed that 55A8 could bind to S proteins from SARS-CoV-2 and the Delta (B.1.617.2), Omicron BA.1 (B.1.1.529.1) and Omicron BA.2 (B.1.1.529.2) variants, as tested by enzyme-linked immunosorbent assay (ELISA) (Fig. 1a). The binding of 58G6, the previously identified NAb mentioned above, was also tested (under review). Furthermore, we found that 55A8 and 58G6 could efficiently recognize other S proteins from a panel of SARS-CoV-2 variants (Extended Data Fig. 1).

**Figure 1.**
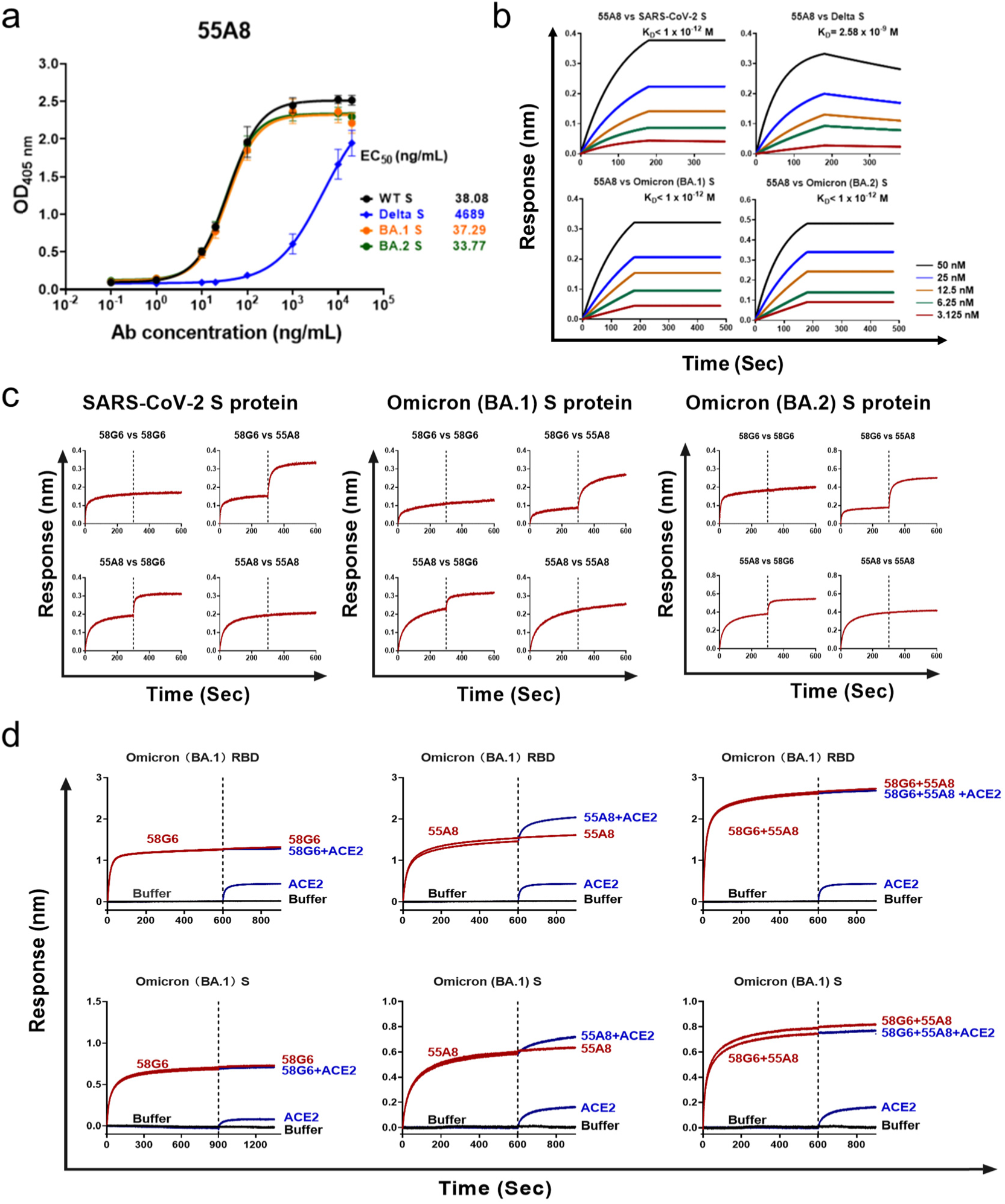
Characterization of 58G6 and 55A8. **A** The binding capabilities of 55A8 against the spike proteins of SARS-CoV-2 and the Delta, Omicron BA.1 and Omicron BA.2 variants were measured by ELISA. **B** The affinities of 55A8 for the S proteins of SARS-CoV-2 and the Delta, Omicron BA.1 and Omicron BA.2 variants were measured by BLI. **C** The sequential binding of 58G6 and 55A8 to the SARS-CoV-2, Omicron BA.1 or Omicron BA.2 S protein measured by BLI. Association was measured first for the antibodies indicated on the left, followed by association of the antibodies indicated on the right. **D** Competition between ACE2 and 58G6, 55A8 or a mixture of 58G6 and 55A8 for binding to the Omicron BA.1 RBD (top) and Omicron BA.1 S protein (down).

The kinetics of 55A8 and 58G6 binding to different S proteins from SARS-CoV-2 and the Delta, Omicron BA.1 and Omicron BA.2 variants were measured by biolayer interferometry (BLI). 55A8 had subpicomolar binding affinities (<10^-12^ M) for Omicron BA.1 and BA.2 (Fig. 1b). In contrast, 58G6 had a lower affinity for Omicron but a high affinity for Delta (under review). Then, we performed BLI to test whether 55A8 and 58G6 could simultaneously bind to the S protein (Fig. 1c). Regardless of which antibody (58G6 or 55A8) first associated with the S protein tested (from SARS-CoV-2, Omicron BA.1 or Omicron BA.2) in the probe, the other antibody could still bind to the S protein noncompetitively.

We also found that 58G6 did compete with ACE2 for binding to the Omicron RBD or S protein, while 55A8 did not compete with ACE2 for binding to the Omicron RBD but partially competed with ACE2 for binding to the Omicron S protein. Interestingly, the combination of 58G6 and 55A8 completely outcompeted ACE2 for binding to the Omicron RBD and S protein (Fig. 1d). Therefore, it will be interesting to test whether the combination of 58G6 and 55A8 could have a synergetic effect.

### 55A8 synergizes with 58G6 for neutralization

To confirm the synergetic effect, we performed a neutralizing assay using pseudoviruses and then authentic viruses. The individual NAbs 55A8 and 58G6 could neutralize pseudotyped SARS-CoV-2 and the Delta, Omicron BA.1 and BA.2 variants, and Omicron BA.1+L452R (L452R might be potential mutation recently predicted in future circulating variants) (Fig. 2a) as well as other SARS-CoV-2 variants (Extended Data Fig. 2). The combination of 58G6 and 55A8 showed obvious synergetic effects against pseudotyped Omicron BA.1, BA.1+L452R, and BA.2 (Fig. 2a). Furthermore, the combination treatment showed synergetic effects against authentic Omicron BA.1 (Fig. 2b). Notably, the IC_50_ for Omicron BA.1 (2.81 ng/mL) was much lower than the IC_50_ values for all other currently known NAbs. In comparison, the IC_50_ values of S309^17^ and LY-CoV1404^18^ are 191.1 and 17.30 ng/mL, respectively. Therefore, the combination of 58G6 and 55A8 exhibited an enhanced neutralizing potency and breadth against SARS-CoV-2.

**Figure 2.**
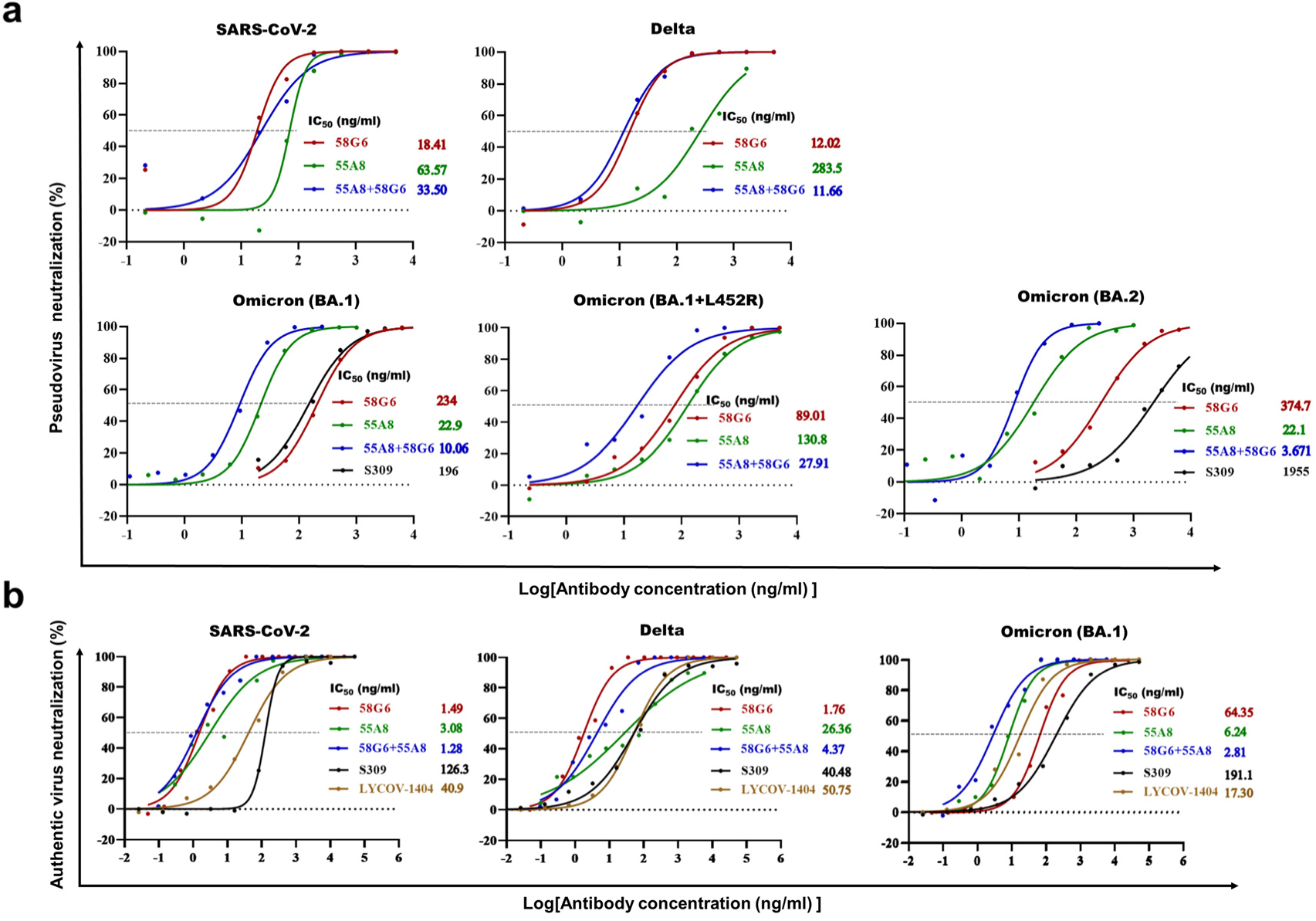
Synergetic effect of 58G6 and 55A8. **a** The neutralizing potencies of 58G6 alone, 55A8 alone or 58G6 and 55A8 in combination against SARS-CoV-2 and the Delta, Omicron BA.1, Omicron BA.1 +L452R, and Omicron BA.2 variants were measured with a pseudovirus neutralization assay. The dashed line indicates a 0% or 50% reduction in viral neutralization. Data are presented as the mean values. **b** Neutralization against authentic SARS-CoV-2, Delta and Omicron BA.1 viruses.

### The 2-antibody cocktail of 58G6 and 55A8 provides synergetic protection *in vivo*

To preliminarily assess whether the combination of 58G6 and 55A8 (2-antibody cocktail, 2-cocktail) could induce synergistic effector function responses *in vivo*, hamsters were intranasally infected with 10^4^ plaque-forming units (PFU) of Omicron BA.1 and treated with 58G6 (1500 µg), 55A8 (500 μg), or 58G6 and 55A8 mixture (2-cocktail, 1000 μg of 58G6 and 300 μg of 55A8) at 1 h pre-infection and 24 and 48 h post-infection (Extended Data Fig. 3a). On day 3 post-infection, the animals were sacrificed, and the turbinates, trachea and lungs were harvested. We observed that all the treatments led to robust viral clearance (Extended Data Fig. 3b and c). Notably, the virus, detected by measurement of the RNA copy numbers and PFU in the upper respiratory tract (turbinate and trachea) and lower respiratory tract (lungs), was completely/or almost completely inhibited in the combination treatment group, while viral RNA or live virus could be detected in the other two groups, which received 58G6 or 55A8 monotherapy. Taken together, these results showed that the combination of 58G6 and 558A could efficiently suppress the Omicron variant *in vivo* through a synergetic effect.

To refine the synergetic effect, more animal protection experiments were performed. To assess the protective efficacy of 55A8 against the Omicron variant, we intranasally challenged hamsters on day 0 with 10^4^ PFU of Omicron BA.1. At 1 h pre-infection and 24 and 48 h post-infection, groups of hamsters received a single intranasal (IN) treatment with 55A8 alone (Group 1, 1000 μg) or 55A8 in combination with 58G6 (Group 3, 500 μg for each antibody). At 3, 24, and 48 h post-infection, groups of hamsters received a single IN treatment with 55A8 alone (Group 2, 1000 μg) or 55A8 in combination with 58G6 (Group 4, 500 μg for each antibody) (Fig. 3a). Another group (Group 5) was treated following the regimen for Group 1 but administered buffer instead of the antibody (Fig. 3a). We measured the effects of 55A8 alone or in combination with 58G6 on Omicron viral replication in clinically relevant tissues (i.e., the nasal turbinates, trachea and lungs), which were collected on day 3 post-infection. Omicron viral replication was detected by RT–qPCR and plaque assays.

**Figure 3.**
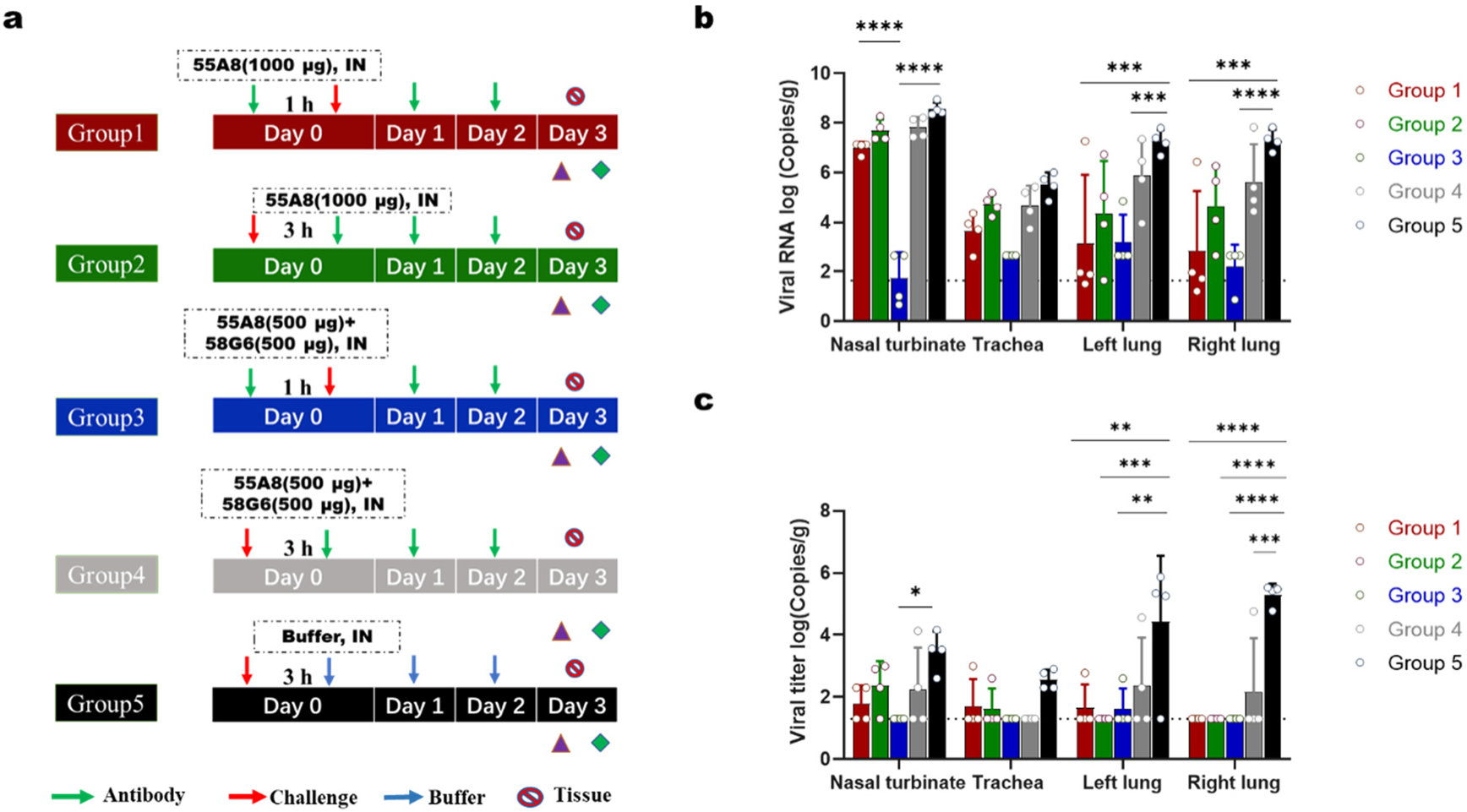
Protective efficacy of 55A8/58G6 cocktails against Omicron in a hamster model. a. Animal experimental scheme. b. Viral RNA (log10(RNA copies per g)) detected in the respiratory tract of hamsters challenged with the Omicron variant at 3 dpi. c. The number of infectious viruses (PFU) in the respiratory tract was measured at 3 dpi with a viral plaque assay performed with Vero E6 cells.

As expected, hamsters treated with buffer had significant viral RNA copy numbers and viral titers in the turbinates and lungs. Pretreatment with 55A8 or the 2-cocktail at 1 h pre-infection followed by two-dose administration lowered the Omicron viral RNA copy number in the lungs by 4–5 logs (Fig. 3a). More notably, in the turbinates, pretreatment with the 55A8 and 58G6 mixture 1 h pre-infection followed by two-dose administration (Group 3) reduced the Omicron viral RNA copy number by six orders of magnitude, producing a copy number that was significantly lower than that in Group 1 (Fig. 3b). Consistently, the viral titers in Group 3 at 3 days post-infection (dpi) were significantly lower than those in the other groups or under the limit the detection (L.O.D.) (Fig. 3c). The protective effects of 55A8 or the 2-cocktail dropped slightly when the treatment was administered at 3 h post-infection and repeated at both 24 and 48 h post-infection. These results indicate that 55A8 protects against IN infection with the Omicron variant and that combining 55A8 with 58G6 increases protective efficiency in hamsters. In fact, the animals treated with the combination of 58G6 and 55A8 maintained a higher weight increase than mock-infected controls (Group 5) (Extended Data Fig. 4). This confirmed that the 55A8/58G6 cocktail reduced Omicron viral replication and prevented disease symptoms without causing additional distress.

To determine the protective doses of the antibodies included in the combination therapy, we further experimented with four different doses (250, 100, 50, and 25 μg for each antibody, administered daily) as Groups 1, 2, 3 and 4, respectively; the antibodies were administered intranasally to evaluate protective efficacy and assess possible differences in protection (Fig. 4a). In addition, another group (Group 5) was administered buffer only as a control. All the tested doses provided protection (Fig. 4b and c). Surprisingly, in the 25 μg dose group (Group 4), the viral RNA level and titer were generally similar to those in the other intervention groups. These data show that the combination of 58G6 and 55A8 confers protection even at the extremely low dose of 25 μg per antibody. Therefore, the 2-cocktail is a promising candidate against Omicron variant infection *in vivo*.

**Figure 4.**
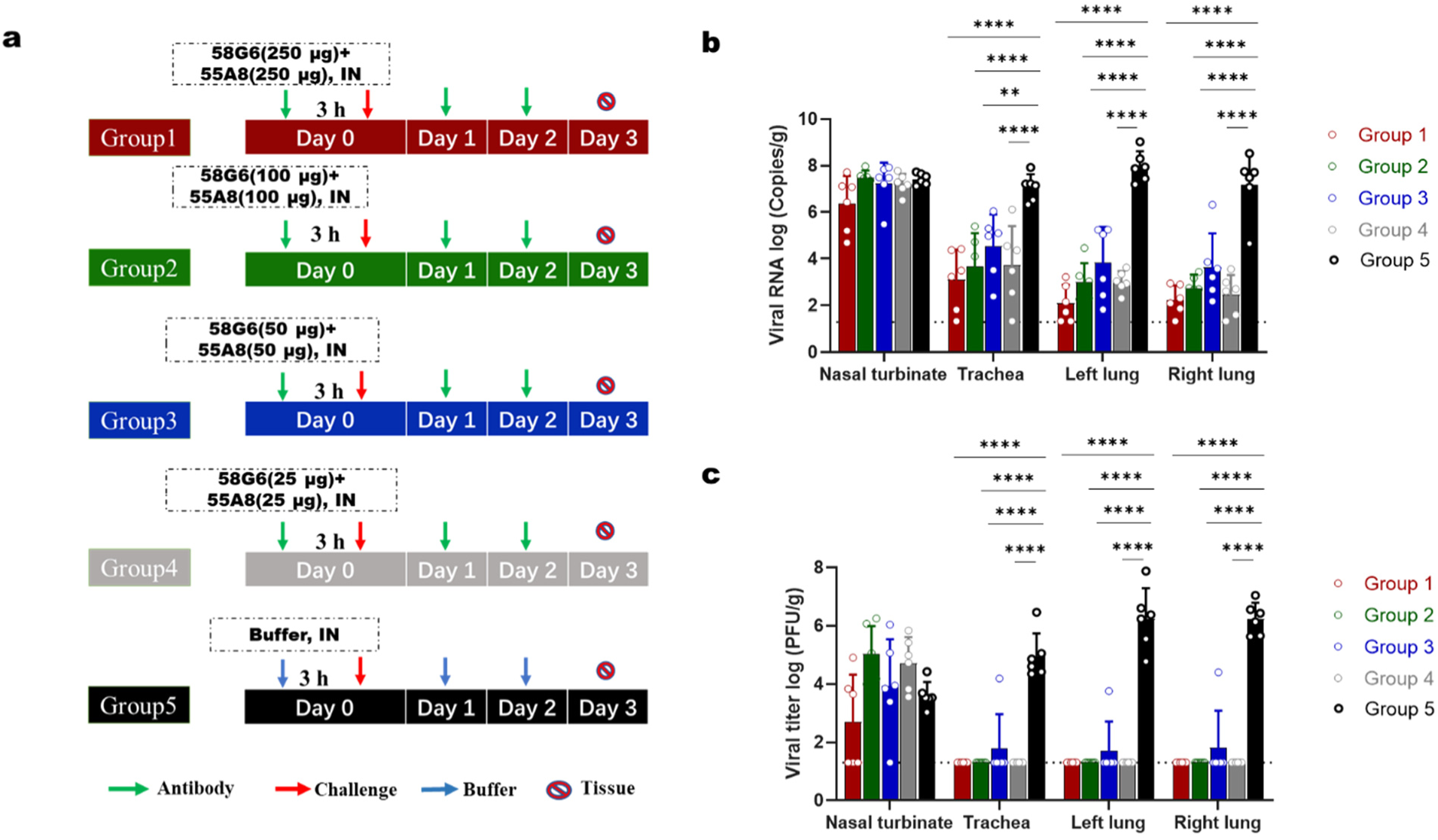
Determination of the prophylactic dose of 55A8/58G6 cocktails against Omicron in the hamster model. **a** Animal experimental scheme. **b** Viral RNA (log10(RNA copies per g)) detected in the respiratory tract of hamsters challenged with the Omicron variant at 3 dpi. **c** The number of infectious viruses (PFU) in the respiratory tract was measured at 3 dpi with a viral plaque assay performed with Vero E6 cells.

### Structure of the complex formed by 55A8 and the S protein

To further investigate the mechanism by which 55A8 neutralizes the Omicron variant, single-particle cryo-electron microscopy (cryo-EM) structures were determined to visualize the antigen-binding fragments (Fabs) of 55A8 in complex with the prefusion Omicron BA.1 S trimer with stabilizing mutations (Extended Data Fig. 6a and c)^19^. When 55A8 Fabs were bound to the Omicron BA.1 S protein, the S trimer adopted a “1-up/2-down” conformation (Class 1) or a “2-up/1-down” conformation (Class 2) (Fig. 5a and b). We refined both to an overall resolution of 3.4 Å (Extended Data Fig. 6e and f), with the majority of selected particle images representing a 3-Fab-per-trimer complex. Further local refinement to 3.3 Å was applied to the same variable domains (including two adjacent up and down RBDs with one 55A8 Fab bound to each RBD) of the two classes, revealing detailed molecular interactions within the binding interface (Extended Data Fig. 6c, Fig. 8a and b). The locally refined density map and the predicted structure of the 55A8 Fab were used to build models to illustrate the structural details of amino acid residues in three dimensions. Superimposition of the up RBDs in the structures of the 55A8 Fab-BA.1 S and ACE2-BA.1 S complexes revealed no overlap between the 55A8 Fab and ACE2 on the same bound RBD, while a steric clash occurred between ACE2 and another 55A8 Fab bound with the adjacent down RBD (Fig. 5c). Considering that 55A8 has a high affinity for the Omicron BA.1 S protein, which is consistent with our cryo-EM observation that most selected particles represent a 3-Fab-per-trimer conformation, the BA.1 RBD-ACE2 interaction can be prohibited by 55A8.

**Figure 5.**
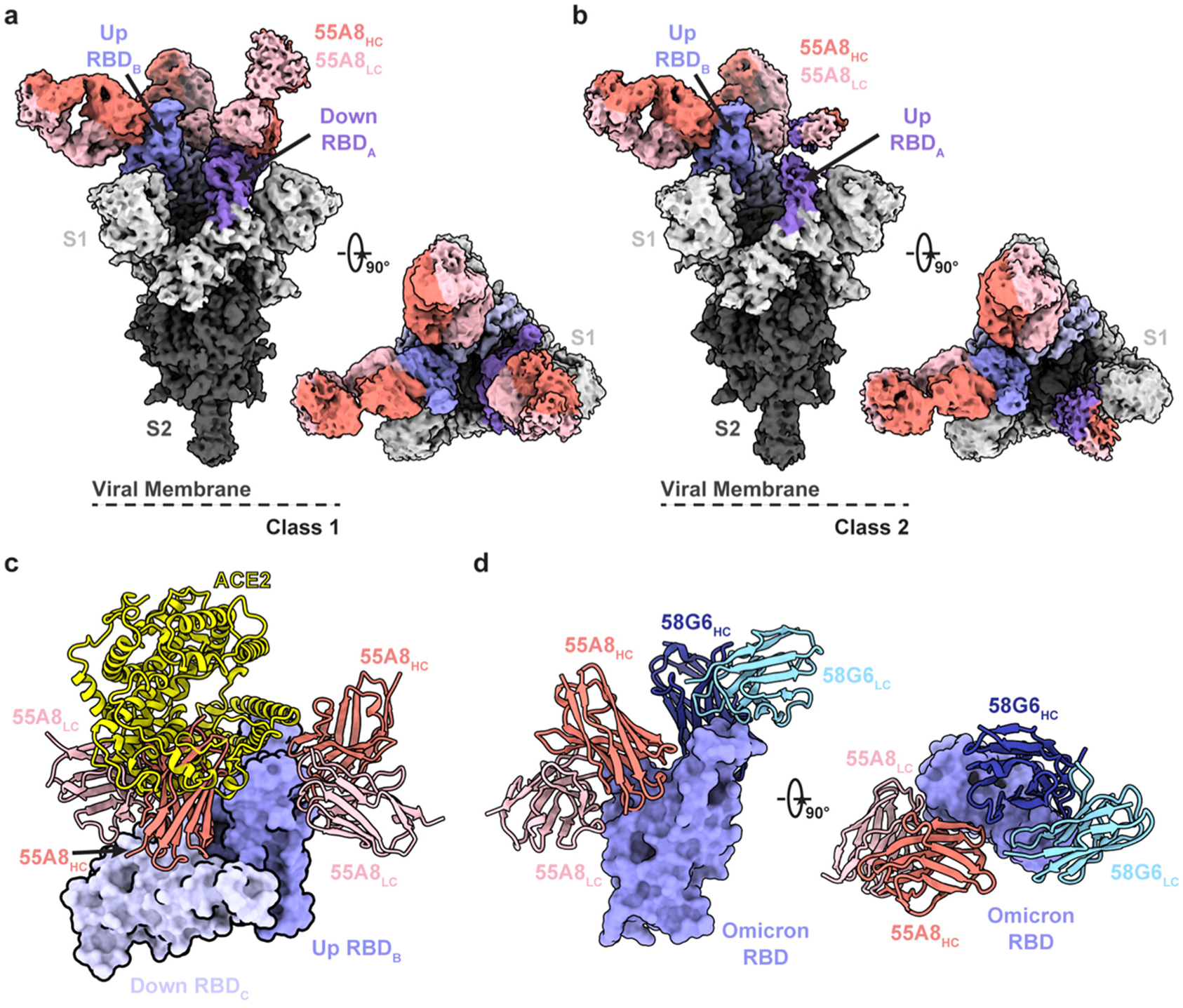
Cryo-EM structures of 55A8 binding to the SARS-CoV-2 Omicron S protein. **a-b** The cryo-EM densities of the 55A8 Fab-Omicron S complex we e observed in two classes (a, Class I, 3.4 Å, 3 Fabs bound with Omicron S in the “2-up/1-down” conformation; b, Class II, 3.4 Å, 3 Fabs bound with Omicron S in the “1-up/2-down” conformation). **c** Superposition of the locally refined Omicron RBD-ACE2 (PDB ID: 7WSA) model together with the locally refined Omicron RBD-55A8 Fab model. **d** Locally refined model of the 55A8 Fab and 58G6 Fab on the same up Omicron RBD. HC, heavy chain; LC, light chain.

Specifically, half of the six complementarity determining regions (CDRs; CDRH3, CDRL1 and CDRL3) of the 55A8 Fab were found to directly participate in the interactions with the S^345 -352^ and S^440 - 450^ regions (Fig. 6a and b). Several potential hydrogen bonds were identified at the interface of the 55A8 Fab and Omicron RBD, representing the unique interaction network between the 55A8 CDRs and the residues within the epitope in the Omicron RBD (Fig. 6c); the sites included R346, Y351, K440, S443, K444, V445 and N450, most of which are conserved among the epidemic variants (except R346K, which was found in the VOC Mu (B.1.621), and N440K, which was found in the VOC Omicron (B.1.1.529))^20^.

**Figure 6.**
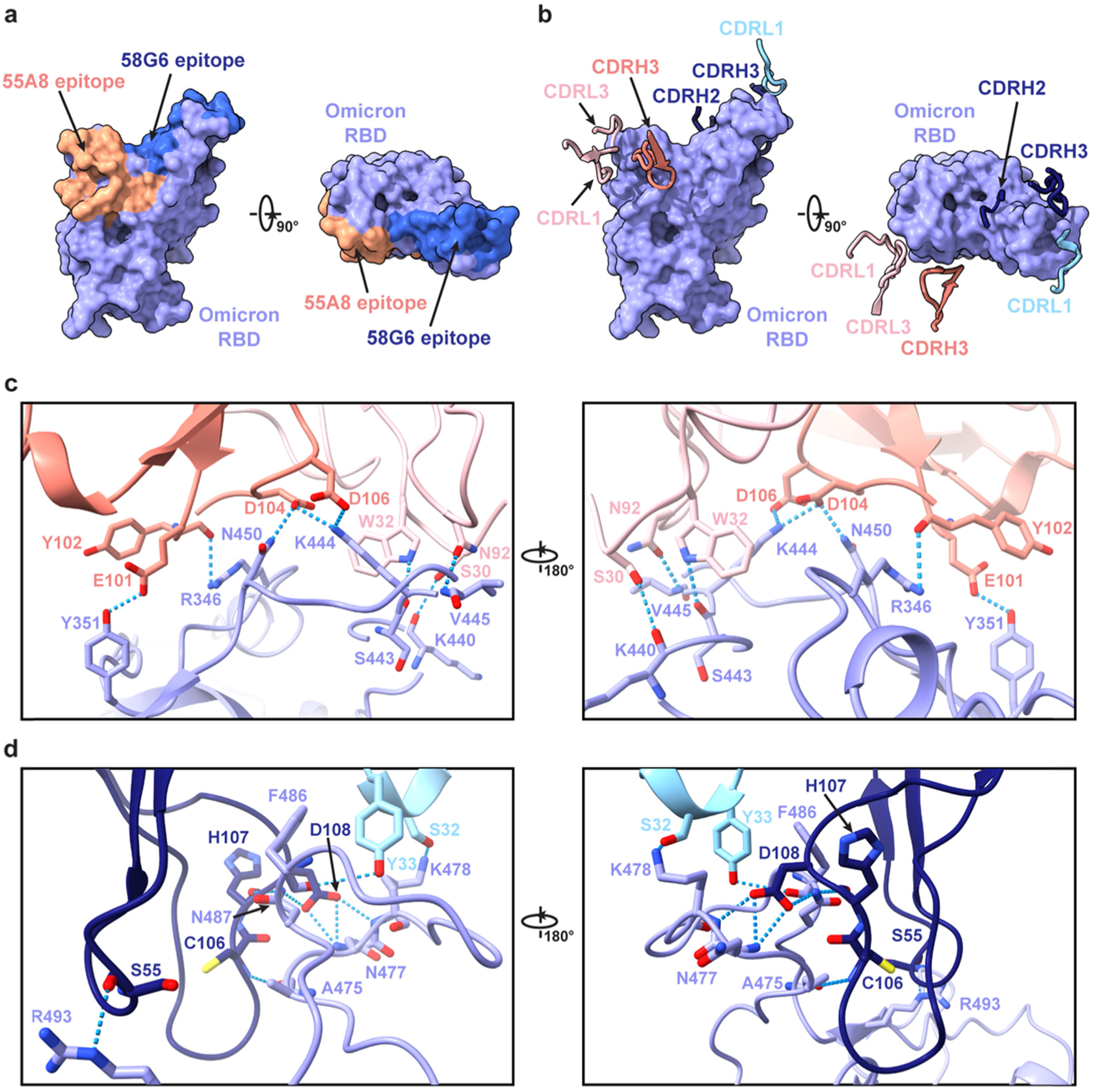
Detailed interactions between 55A8/58G6 cocktails and the Omicron RBD. **A** Surface representation of the 55A8 and 58G6 epitopes on the Omicron RBD surface. **b** CDR loops of the 55A8 Fab and 58G6 Fab overlaid on the surface representation of the Omicron RBD. **c, d** Potential hydrogen bonds (shown as blue dashed lines) at the binding interface between the Omicron RBD and 55A8 Fab (c) or 58G6 Fab (d).

### Synergetic mechanism of the 2-cocktail

The combined use of several noncompeting antibodies, or cocktails, could be an effective treatment to reduce selective pressure and prevent immune escape mutations. Another previously identified SARS-CoV-2 NAb, 58G6, also showed neutralizing potency against the WT virus. 58G6 recognizes the S^450 - 458^ and S^470 - 495^ regions in the WT RBD, while 55A8 binds to S^345 - 352^ and S^440 - 450^ in the Omicron RBD. The epitopes recognized by the two antibodies are located in different regions of the RBD, suggesting the potential for combined use of 58G6 and 55A8 against the Omicron variant.

To further investigate the molecular mechanism underlying the effect of the 2-cocktail, we determined the cryo-EM structures of the Omicron BA.1 S trimer in complex with the 55A8 and 58G6 Fabs (Extended Data Fig. 6b and d). Consistent with Class 1 and Class 2 containing 55A8 alone (Extended Data Fig. 7a), the BA.1 S trimer adopted a “1-up/2-down” conformation (Class 3) or a “2-up/1-down” conformation (Class 4) when treated with 55A8 and 58G6 (Extended Data Fig. 7b and d). We refined both to an overall resolution of 3.3 Å (Extended Data Fig. 6g and h), with the majority of the selected particle images representing a 4-Fab-per-trimer complex. The 55A8 Fab and 58G6 Fab simultaneously bound to a single up RBD, exhibiting no conformational changes compared with the BA.1 S-55A8 Fab complex. 58G6 could occupy the ACE2 binding site on the Omicron BA.1 RBD and further occlude the accessibility of the Omicron BA.1 S protein to ACE2 (Fig. 5d). Superimposition of the WT RBD-58G6 structure model (PDB ID: 7E3L) and the locally refined BA.1 RBD-58G6 model revealed no significant conformational changes, suggesting that 58G6 utilizes the same binding mode to bind to the WT and Omicron BA.1 S proteins (Fig. 6d, Extended Data Fig. 8c and d).

## Discussion

To date, the COVID-19 pandemic continues. To combat SARS-CoV-2, dozens of vaccines, NAbs and small-molecule drugs have been approved for the prevention and treatment of SARS-CoV-2 infection. For example, approximately ten vaccines are currently approved for use by the World Health Organization (WHO)^21^. However, breakthrough infections in fully vaccinated individuals have been reported worldwide, which indicates that the approved vaccines might primarily decrease mortality rather than prevent infection^22–25^. Hence, more potent antiviral agents that could provide preventive protection, such as NAbs, are still urgently needed.

In November 2021, the Omicron variant (B.1.1.529) was first detected in Gauteng Province, South Africa^26^. Compared to SARS-CoV-2, Omicron BA.1 has 15 mutations in the RBD, while BA.2 has 16 mutations. BA.1 and BA.2 appear to be very resistant to Class I, II and IV NAbs while very small part of Class III NAbs could still efficiently neutralize BA.1 and/or BA.2^4^. Our antibody 55A8 is a broad NAb that potently neutralizes SARS-CoV-2 and Omicron BA.1 and BA.2 without significant differences. Notably, its activity is comparable to or even better than that of other NAbs (e.g., LY-CoV1404) that have been reported to retain neutralizing potency against the Omicron variant^27–32^.

Antibody cocktails have been shown to be outstanding in the control of viral infections, as they may have relatively good antiviral potency and resistance to mutations. The first use of an antibody cocktail strategy (e.g., ZMapp^33^) in patients was in the treatment of infections caused by Ebola virus (EBOV). Following this strategy, REGN-EB3 (Inmazeb), a three-NAb cocktail, was approved by the U.S. Food and Drug Administration (FDA) as a treatment for EBOV infection^34^. Currently, several cocktails, such as casirivimab/imdevimab^35^, bamlanivimab/etesevimab^36^, and amubarvimab/romlusevimab^37^, have also been successfully authorized to prevent and treat SARS-CoV-2 infection. Unfortunately, the activity of these cocktails is dramatically challenged by Omicron, especially the sublineage BA.2. Therefore, the EUAs for most of these cocktails have been cancelled. To date, no authorized antibody cocktail can neutralize both BA.1 and BA.2.

Based on the BLI competition experiment and cryo-EM, 58G6 and 55A8 were concluded to have different binding sites. 55A8 recognized sites localized on the outer face of the RBD, but 58G6 mainly bound to the RBM. The two monoclonal antibodies (mAbs) were shown to bind to nonoverlapping epitopes in the RBD and block RBD binding with ACE2 in different ways. We noticed that 55A8 exhibited slightly decreased neutralizing potency against VOC Delta, Lambda and Kappa, which might be due to the mutation at L452 site. Delta and Kappa carry the mutation L452R, and Lambda carries the mutation L452Q. 58G6 exhibited slightly decreased neutralizing potency against the Omicron BA.1 and BA.2 variants. Therefore, we combined 55A8 with 58G6 to create a complementary cocktail (2-cocktail). Importantly, the 2-cocktail potently neutralized SARS-CoV-2, Delta, Omicron BA.1 and Omicron BA.2 pseudoviruses as well as authentic SARS-CoV-2, Delta, and Omicron BA.1 viruses.

We used cryo-EM to elucidate the detailed interactions between 55A8 and Omicron BA.1 S trimers. The binding sites of 55A8 were found to be in the S^345 - 352^ and S^440–450^ regions, not in the RBM, and these regions are more conserved (except N440K) than the RBM. This also explained why the mAb 55A8 still showed extremely high affinities for the Omicron BA.1 and BA.2 S trimers. Superimposition of the up RBD in the structures of the 55A8 Fab-BA.1 S and ACE2-BA.1 S complexes revealed no overlap between the 55A8 Fab and ACE2 on the same bound RBD, which explained why the mAb 55A8 was not able to compete with ACE2 for Omicron RBD binding.

When 55A8 Fabs were bound to BA.1 S trimers, the S trimer adopted a “1-up/2-down” conformation or a “2-up/1-down” conformation. In the “1-up/2-down” conformation, two 55A8 Fabs were bound to two down RBDs and occluded ACE2 binding to the remaining up RBD. In the “2-up/1-down” conformation, one 55A8 Fab bound to one down RBD and could occlude ACE2 binding to an adjacent up RBD while freeing the other up RBD for binding to ACE2, which explains the partial competition between 55A8 and ACE2 for Omicron BA.1 S trimer binding.

To further evaluate the potential of the combination of 58G6 and 55A8, we determined the cryo-EM structures of Omicron BA.1 S trimers in complex with 55A8 and 58G6 Fabs. Consistent with using 55A8 alone, the BA.1 S trimer also adopted a “1-up/2-down” conformation or a “2-up/1-down” conformation, both of which represent a 4-Fab-per-trimer complex with one 58G6 Fab binding to one up RBD and three 55A8 Fabs binding to all three RBDs. 58G6 can occupy the ACE2 binding site in the up RBD and further occlude the accessibility of BA.1 S to ACE2, which completely outcompetes ACE2 and explains the synergetic effects of 55A8 and 58G6.

58G6 is an RBM-targeted NAbs with retained neutralizing potency against Omicron BA.1 and BA.2. In our previous structure of 58G6 bound to WT S, we found that 58G6 could bind to two epitopes in S^450–458^ and S^470–495^ at the same time. Omicron does not have mutations in the S^450–458^ region but does have four mutations in the S^470–495^ region, S477N, T478K, E484A, and Q493R. 58G6 can form hydrogen bonds with the mutated amino acids N477, K478 and R493, suggesting that this NAb has retained neutralizing potency against Omicron compared with other RBM-targeted NAbs. This unique epitope recognized by 58G6 enables this NAb to be used in cocktails with Class III antibodies, which include most NAbs that retain neutralizing potency against Omicron.

Our combination therapy including 55A8 and 58G6 (2-cocktail) showed great synergetic effects against Omicron infection *in vitro* and *in vivo*. Notably, the synergetic effect dramatically reduces the dose of the cocktail needed to control infection according to the animal study, which indicates that the cocktail has the potential to relieve the disease burden in patients. In addition, because the cocktail can be delivered nasally, it will be safer and more acceptable to different patients and can be easily distributed. Hence, this 2-cocktail is not only a good candidate for combating currently circulating SARS-CoV-2 variants but also a promising drug candidate for use against future SARS-CoV-2 variants. The unique epitope and neutralizing mechanism disclosed here could also provide important clues for the development of more promising vaccines and NAbs.

## Methods

### Ethics statements

All procedures associated with the animal study were reviewed and approved by the Institutional Animal Care and Use Committee of the Institute of Wuhan Institute of Virology, Chinese Academy of Sciences, and were performed in an ABSL-3 facility (ethical approval number: WIVA45202104).

### Cells and viruses

The SARS-CoV-2 WIV04 strain (IVCAS 6.7512) was originally isolated from a COVID-19 patient in 2019 (GISAID, accession no. EPI_ISL_402124)^38^. The SARS-CoV-2 Delta variant (B.1.617.2; GWHBEBW01000000) was kindly provided by Hongping Wei’s laboratory, Wuhan Institute of Virology, Chinese Academy of Science. The SARS-CoV-2 Omicron virus was isolated from a throat swab of a patient in Hong Kong by the Institute of Laboratory Animal Sciences, Chinese Academy of Medical Sciences (CCPM-B-V-049-2112-18). Viral stocks were prepared by propagation in Vero E6 cells (catalog no. ATCC® CRL-1586™) in Dulbecco’s modified Eagle’s medium (DMEM) supplemented with 10% fetal bovine serum (FBS) and 1% penicillin and streptomycin (P/S). All the cells were cultured under 5% CO_2_ in a humidified incubator at 37°C. All live virus experiments were performed at the Wuhan Institute of Virology in a biosafety level 3 (BSL3) containment facility under an approved biosafety use authorization.

### Animals

Female Syrian hamsters (five to six weeks of age) were purchased from Wuhan Institute of Biological Products Co., Ltd.

### Protein expression and purification

The expression and purification procedures for the SARS-CoV-2 Omicron S protein were described previously^39^. In brief, the gene encoding stabilized Omicron S ECD HexaPro was constructed and expressed using FreeStyle 293-F cells. Protein was purified from filtered cell supernatants using Ni Sepharose resin (GE Healthcare, USA) before being subjected to additional purification by gel filtration chromatography using a Superose 6 10/300 increase column (GE Healthcare, USA) in 1× phosphate-buffered saline (PBS, pH 7.4).

All the antibodies were prepared as previously described^16^. In brief, two plasmids containing the light chain and heavy chain of antibodies were transiently cotransfected into Expi293™ cells (ThermoFisher, A14528) at a 1:1 ratio with ExpiFectamine™ 293. After 7 days, antibodies were purified from filtered cell supernatants using protein G Sepharose (GE Healthcare) and dialyzed into 1× PBS (pH 7.4)

### ELISA

384 wells ELISA plates (NUNC Clear Flat-Bottom Immuno Nonsterile 384-Well Plates, Cat no. 464718) were coated with 10 μL/well of S proteins from SARS-CoV-2 WT and different variants (Sino Biological, Beijing, China) at 2 μg/mL in PBS, pH 7.4, 4°C overnight. Serially diluted 55A8 or 58G6 solution was added to each well with 10 μL/well and incubated at 37°C for 1h. ALP-conjugated anti-human IgG antibody (ThermoFisher, A18808, 1:2000) was used as the detection antibody at 37°C for 30 min . For the quantification of bound IgG, PNPP (ThermoFisher, 34045) was added and the absorbance was measured at 405 nm using MultiSkan GO fluoro-microplate reader (ThermoFisher).

### Biolayer interferometry (BLI)

Bio-layer interferometry assays were conducted on Octet R2 Protein Analysis System (Fortebio). Protein biotinylation was performed using the EZ-link NHS-PEO Solid Phase Biotinylation Kit (Pierce) and purified using MINI Dialysis Unit (ThermoFisher). For measurement of the affinities of 55A8 and 58G6 for SARS-CoV-2 and the Delta, Omicron BA.1 and Omicron BA.2 variants, streptavidin (SA) biosensors (ForteBio) were immersed with biotinylated mAbs to generate capture mAbs, and then the sensors were immersed in buffer (0.02% Tween-20, 1 mg/mL BSA in PBS) to the baseline. After association with twofold diluted proteins (diluted from 50 nM to 3.125 nM), disassociation was conducted.

For the mAb competition experiments, biotinylated S proteins were loaded at 1 nm onto SA biosensors, and mAb association was performed at 20 μg/mL for 300 s. For ACE2 competition experiments, the biotinylated RBD and S were loaded at 1 nm and 3 nm, respectively, at 1 μg/mL onto SA biosensors. The first antibody was allowed to associate for 600 s at 20 μg/mL, and the second protein (ACE2 (50 μg/mL) or a mixture of equal amounts of antibodies and ACE2) was allowed to associate for 300 s.

### Inhibition of pseudotyped SARS-CoV-2 infection

These experiments were performed as described previously with a minor change^16^. In brief, serially diluted mAbs in a volume of 50 μl were incubated with the same volume of Lenti-X293T cell supernatants containing pseudovirus (SARS-CoV-2, Delta, Omicron BA.1 or Omicron BA.2) for 1 h at 37°C. These pseudovirus/antibody mixtures were added to ACE2-expressing Lenti-X293 T cells (293 T/ACE2 cells). After 8 h, the supernatants were replaced with fresh culture medium. After 24 h, the luciferase activities of infected 293T/ACE2 cells were detected with the Bright-Luciferase Reporter Assay System (Beyotime, RG055M). The IC_50_ values of the evaluated mAbs were determined with a Varioskan LUX Microplate Spectrophotometer (ThermoFisher) and calculated by four-parameter logistic regression using GraphPad Prism 8.0.

### Neutralizing activity against authentic SARS-CoV-2

Neutralization titers of antibodies were measured with a plaque reduction neutralization test (PRNT) using authentic SARS-CoV-2, Delta, and Omicron viruses^40^. Briefly, Vero E6 cells (2.5 × 10^5^) were seeded in each well of 24-well culture plates. The cells were incubated overnight at 37°C with 5% CO_2_. On the following day, each antibody was serially diluted 5-fold in the culture medium with the highest concentration being 100 μg/ml. The diluted antibody was incubated with an equal volume including 600 PFU/ml SARS-CoV-2 at 37°C for 1 h, after which the antibody-virus mixtures were inoculated onto preseeded Vero E6 cell monolayers in 24-well plates. After 1 h of infection, the inoculum was removed, and 100 μl of overlay medium (DMEM supplemented with 0.8% methylcellulose, 2% FBS, and 1% P/S) was added to each well. After incubating the plates at 37°C for 96 h, the plates were fixed with 8% paraformaldehyde and stained with 0.5% crystal violet. The plaques in each well were counted and normalized to the non-antibody-treated controls to calculate relative infectivity. PRNT_50_ values were calculated in GraphPad Prism 9.

### Animal protection experiments

To assess whether the 58G6 and 55A8 antibody cocktail induced synergistic effector function responses *in vivo*, hamsters were infected with 10^4^ PFU of Omicron and treated with 58G6 (1500 μg), 55A8 (500 μg), or the 58G6 and 55A8 mixture (1000 μg of 58G6 and 300 μg of 55A8) at 1 h pre-infection and 24 and 48 h post-infection. On day 3 post-infection, the animals were sacrificed, and the turbinates, trachea and lungs were harvested. To compare the protective efficiency of 55A8 and the 2-cocktail, another animal protection experiment was performed. Syrian golden hamsters were infected through IN inoculation of 10^4^ PFU of Omicron variant in 100 μl of DMEM per animal after anesthetization with isoflurane. At 1 h pre-infection, animals were given an IN dose of 1000 μg of 55A8 (Group 1, n =4) or 500 μg of 55A8 and 500 μg of 58G6 (Group 3, n =4). Then, doses (1000 μg 55A8 or the antibody cocktail containing 500 μg of 55A8 and 500 μg of 58G6) were administered on days 1 and 2 post-infection in Group 1 and Group 3, respectively. At 3 h post-infection, animals were given an IN dose of 1000 μg of 55A8 (Group 2, n =4) or the two-antibody cocktail (Group 4, n =4). On days 1 and 2 post-infection, doses (1000 μg of 55A8 or the antibody cocktail) were administered in Group 2 and Group 4, respectively. The hamsters were then weighed daily for the duration of the study. On day 3 post-infection, the animals were sacrificed, and the turbinates, trachea and lungs were harvested. For the infected control experiment, 6-week-old Syrian golden hamsters were intranasally treated with buffer containing either 55A8 or 58G6, and the turbinates, trachea and lungs were harvested at three dpi. Lung tissue was analyzed by viral titering (by PFU/g and qRT–PCR) and histological evaluation.

To determine the antibody cocktail dose that could provide protection, we performed additional animal experiments. A range of doses (500 to 50 μg) of the two-antibody cocktail were inoculated into hamsters intranasally. Three hours later, the golden hamsters were anesthetized with isoflurane and intranasally inoculated with 10^4^ PFU of Omicron variant. Then, two doses of the antibody cocktail were administered at 24 h and 48 h. The animals were monitored daily, and weight was measured at the indicated time points. On day 3 post-infection, the animals were sacrificed, and the turbinates, trachea, and lungs were harvested. Lung tissue was analyzed by viral titering (by PFU/g and qRT–PCR) and histological evaluation.

### RT–qPCR assay

To determine the RNA copy numbers in different tissues, SARS-CoV-2 genomic mRNA was assessed by RT–qPCR as previously described^41^ using the following primer and probe sequences: forward, ORF1ab-F, 5’-CCCTGTGGGTTTTACACTTAA-3’ and ORF1ab-R, 5’-ACGATTGTGCATCAGCTGA-3’.

### Plaque assay for SARS-CoV-2

The viral titers in different tissues were determined with a plaque assay performed as previously described with slight modification^42^. Briefly, virus samples were 10-fold serially diluted in culture medium (first dilution, 1:5) and inoculated onto Vero E6 cells seeded overnight at 1.5 × 10^5^/well in 24-well plates. After a 1-h incubation at 37°C, the inoculum was replaced with DMEM containing 2.5% FBS and 0.9% carboxymethyl-cellulose. The plates were fixed with 8% paraformaldehyde and stained with 0.5% crystal violet 4 days later.

### Histological analysis

Paraffin-embedded hamster lung tissue blocks were cut into 5-mm sections. The sections were stained with hematoxylin and eosin (H&E) and analyzed. H&E-stained sections of hamster lungs were scored according to a standardized scoring system evaluating the presence of interstitial pneumonia, type II pneumocyte hyperplasia, edema, fibrin deposition, and perivascular lymphoid cuffing^43^.

### Cryo-EM sample preparation and data collection

To prepare the complex formed by 55A8 and the SARS-CoV-2 Omicron S protein, purified Omicron S was diluted to a concentration of 2.0 mg/mL in PBS (pH 7.4) and incubated with the 55A8 Fab at a molar ratio of 1:5. The mixture was incubated on ice for 1 h and then subjected to gel filtration chromatography using a Superose 6 10/300 column (GE Healthcare, USA) in 1× PBS (pH 7.4). The complex sample was concentrated to 0.8 mg/mL, and a 2.5-μL aliquot of the mixture was applied to an H_2_/O_2_ glow-discharged, 300-mesh Quantifoil R1.2/1.3 copper grid (Quantifoil, Micro Tools GmbH, Germany). The grid was then blotted for 2.5 s with a blot force of -1 at 8°C and 100% humidity and plunge-frozen in liquid ethane using a Vitrobot (Thermo Fisher Scientific, USA).

To prepare the complex formed by 55A8, 58G6 and the SARS-CoV-2 Omicron S protein, purified Omicron S was diluted to a concentration of 0.8 mg/mL in PBS (pH 7.4). Ten microliters of purified SARS-CoV-2 S was mixed with 0.6 μL of 5 mg/mL 58G6 Fab fragments in PBS and incubated for 15 minutes on ice. Then, 0.7 μL of 4 mg/mL 55A8 Fab fragments in PBS was applied and incubated for 15 minutes on ice. A 3 μL aliquot of the mixture (added with 0.01% DDM) was applied to an H_2_/O_2_ glow-discharged, 300-mesh Quantifoil R1.2/1.3 grid (Quantifoil, Micro Tools GmbH, Germany). The grid was then blotted for 3.0 s with a blot force of -1 at 8 °C and 100% humidity and plunge-frozen in liquid ethane using a Vitrobot (Thermo Fisher Scientific, USA).

Cryo-EM datasets were collected on a 300 kV Titan Krios microscope (Thermo Fisher Scientific, USA) equipped with a K3 detector (Gatan, USA). The exposure time was set to 2.4 s with a total accumulated dose of 60 electrons per Å^2^, which yielded a final pixel size of 0.82 Å. A total of 1,525 micrographs were collected for the Omicron S-55A8 Fab complex in a single session with a defocus range between 1.2 and 2.2 μm using SerialEM^44^, while 3,605 micrographs were collected for the Omicron S-55A8 Fab-58G6 Fab complex under conditions similar to those described previously. The statistics for cryo-EM data collection can be found in Extended Data Tables 1 and 2.

### Cryo-EM data processing

All dose-fractioned images were motion-corrected and dose-weighted with MotionCorr2 software^45^, and their contrast transfer functions (CTFs) were estimated by cryoSPARC patch CTF estimation^46^. For the dataset for the Omicron S-55A8 Fab complex, a total of 847,678 particles were autopicked, and after 2D classification, 262,990 particles were selected for ab-initio reconstruction in six classes. These six classes were used as 3D volume templates for heterogeneous refinement with all selected particles, with 181,072 particles converged into the Omicron S triple-bound 55A8 class. Next, this particle set was used to perform ab-initio reconstruction again in four classes. After heterogeneous refinement with all selected particles, 82,383 particles converged into Class 1 (“1-up/2-down” conformation), and 94,894 particles converged into Class 2 (“2-up/1-down” conformation). Next, these two particle sets were used to perform nonuniform refinement, yielding a resolution of 3.4 Å for both triple-bound Omicron S-55A8 Fab complexes.

For the dataset for the Omicron S-55A8/58G6 Fab complex, a total of 914,631 particles were autopicked, and after 2D classification, 608,282 particles were selected for several rounds of ab-initio reconstruction in six classes. These six classes were used as 3D volume templates for heterogeneous refinement with all selected particles. Finally, 216,663 particles converged into Class 3 (“1-up/2-down” conformation), and 204,153 particles converged into Class 4 (“2-up/1-down” conformation). Next, these two particle sets were used to perform nonuniform refinement, yielding a resolution of 3.3 Å for both triple-bound Omicron S-55A8 Fab complexes.

To improve the local resolution at the binding interface, we subsequently added local refinement processing. A local reconstruction focusing on two adjacent up and down RBDs with one 55A8 Fab bound to each RBD was carried out. Furthermore, the density map for the binding interface could be further improved by local averaging of the RBD-55A8 Fab equivalent copies, ultimately yielding a 3.3-Å map of the region corresponding to the RBD-55A8 Fab interface. Similarly, we improved the local resolution between the 55A8 and 58G6 variable domains and the RBD up to 3.3 Å.

Local resolution estimation, filtering, and sharpening were also carried out using cryoSPARC. The full cryo-EM data processing workflow is described in Extended Data Fig. 6, and the model refinement statistics can be found in Extended Data Tables 1 and 2.

### Model building and refinement

To build the structures of the Omicron S-55A8 Fab complex, the structure of the Omicron S-510A5 complex model (PDB ID: 7WS5)^39^ was placed and rigid-body fitted into cryo-EM electron density maps using UCSF Chimera^47^. The 55A8 Fab model was first predicted using Phyre2^48^ and then manually built in Coot 0.9^49^ with the guidance of the cryo-EM electron density maps, and overall real-space refinements were performed using Phenix 1.19^50^. The data validation statistics are shown in Extended Data Table 1.

To build the structures of the Omicron S-55A8/58G6 Fab complex, the previously described structure of the Omicron S-55A8 complex model was placed and rigid-body fitted into cryo-EM electron density maps using UCSF Chimera. The 58G6 Fab model^15^ was manually built in Coot 0.9 with the guidance of the cryo-EM electron density maps, and overall real-space refinements were performed using Phenix 1.19. The data validation statistics are shown in Extended Data Table 2.

### Statistical analysis

Statistical analyses were performed using GraphPad Prism software v.9.2.0. Comparisons between two groups were performed using unpaired Student’s t tests. Comparisons among multiple groups were performed using one-way ANOVA followed by Tukey’s multiple comparison post hoc test. P < 0.05 was considered significant (significance is denoted as follows: *P < 0.05, **P < 0.01, ***P < 0.001, and ****P < 0.0001).

## Data availability

The coordinates and cryo-EM map files for the 55A8-BA.1 S complexes and 55A8/58G6-BA.1 S complexes have been deposited in the Protein Data Bank (PDB) under accession number 7WWI, 7WWJ, 7WWK, 7XJ6, 7XJ8 and 7XJ9. All other data are available from the corresponding author upon reasonable request.

## Acknowledgments

We thank the Center for Animal Experiment and BSL-3 laboratory, Wuhan Institute of Virology, Chinese Academy of Sciences; Center for Biosafety Mega-Science, Chinese Academy of Sciences; and the National Virus Resource Center for resource support. This work was jointly supported by the Natural Science Foundation of Hubei Province of China (2019CFA076); the National Natural Science Foundation of China (32170949, 81871639, 92169109, 81871656 and 8181101099); the National Science and Technology Major Project (2017ZX10202203); the Emergency Project from the Science & Technology Commission of Chongqing (cstc2021jscx-fyzxX0001); the National Key Research and Development Program of China (2018YFA0507100 and 2016YFD0500300); Lingang Laboratory (LG202101-01-07); and Science and Technology Commission of Shanghai Municipality (YDZX20213100001556 and 20XD1422900).

## Author Contributions

Ailong Huang, Sandra Chiu, Aishun Jin and Haitao Yang conceived and designed the project. For biological function analysis of the NAbs, Feiyang Luo, Tingting Li, Meiying Shen, Xiaojian Han, Yingming Wang and Chao Hu screened and cloned the antibodies, and expressed and purified the antibodies; Feiyang Luo, Tingting Li and Wei Wang were responsible for BLI assays for the binding ability, the affinity and the competition experiment of NAbs; Feiyang Luo, Tingting Li, Shuyi Song, Kai Wang and Ni Tang prepared various pseudovirus and conducted the pseudovirus neutralization assays. For the efficacy test of the NAbs *in vitro* and *in vivo*, Xinghai Zhang, Huajun Zhang, Junhui Zhou, Shaohong Chen, Yan Wu and Rui Gong performed authentic SARS-CoV-2 neutralization assays and animal experiments. For structure analysis, Hangtian Guo and Yuchi Lu cloned, expressed and purified Omicron BA.1 S proteins; Hangtian Guo, Yan Gao, and Yuchi Lu collected, processed the cryo-EM data, and built and refined the structure model; Xiaoyun Ji, Haitao Yang and Tengchuan Jin analyzed and discussed the cryo-EM data. Ailong Huang, Sandra Chiu, Aishun Jin, Xiaoyun Ji, Rui Gong and Xinghai Zhang wrote the manuscript. All authors revised and reviewed the final manuscript.

---

**Extended Data Figure 1.**
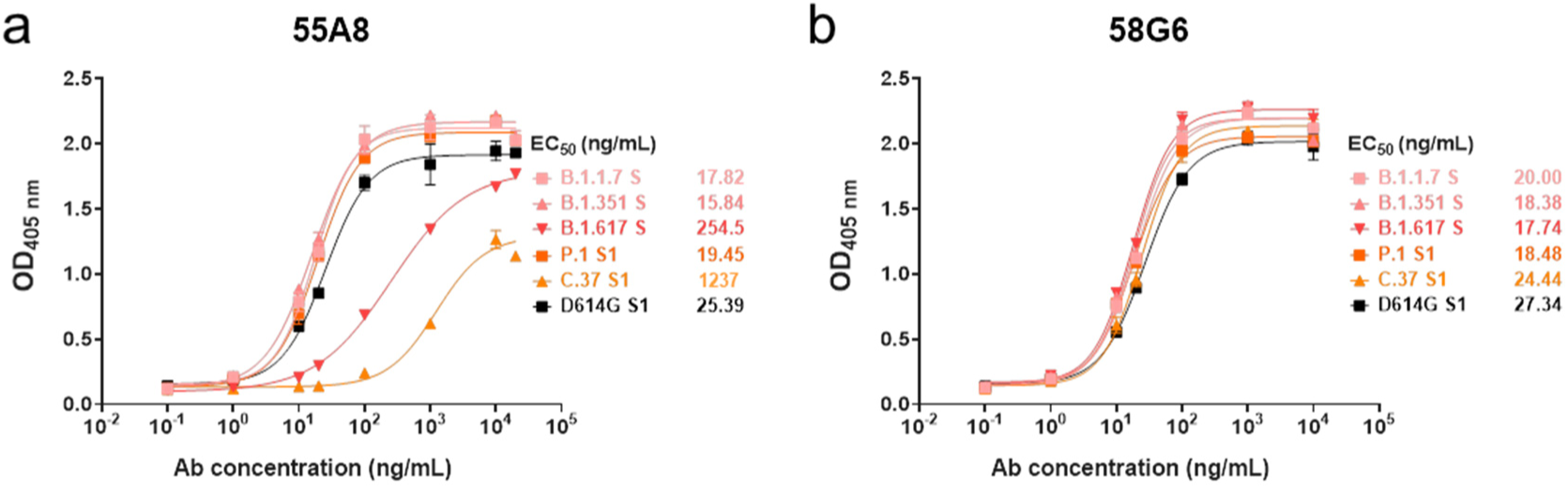
The binding capabilities of 55A8 and 58G6 against the S proteins of several SARS-CoV-2 variants. **a-b** The binding capabilities of 5A8 and 58G6 against the S proteins of SARS-CoV-2 and the strains B.1.1.7, B.1.351, B.1.617, P.1, C.37 and D614G were measured by ELISA.

**Extended Data Figure 2.**
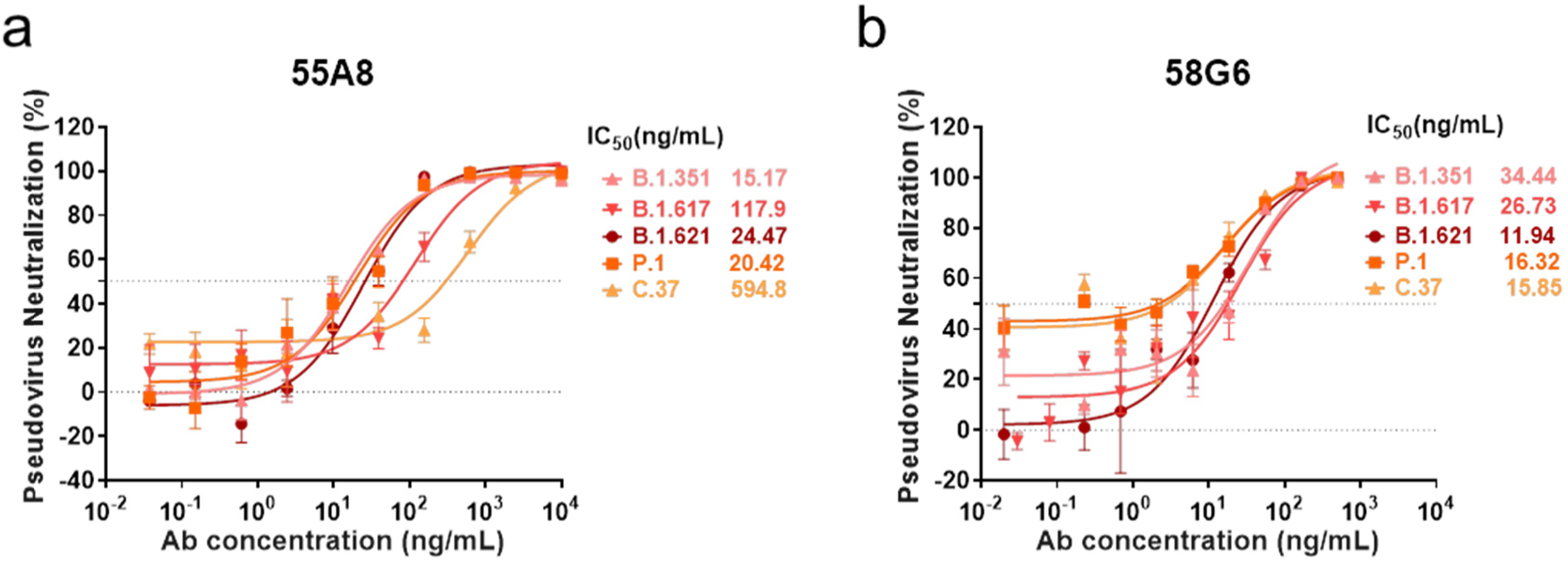
The neutralizing potencies of 55A8 and 58G6 against several SARS-CoV-2 variant pseudoviruses. a-b. The neutralizing potencies of 55A8 and 58G6 against the variants B.1.1.7, B.1.351, B.1.617, B.1.621, P.1, C.37 and D614G were measured with a pseudovirus neutralization assay.

**Extended Data Figure 3.**
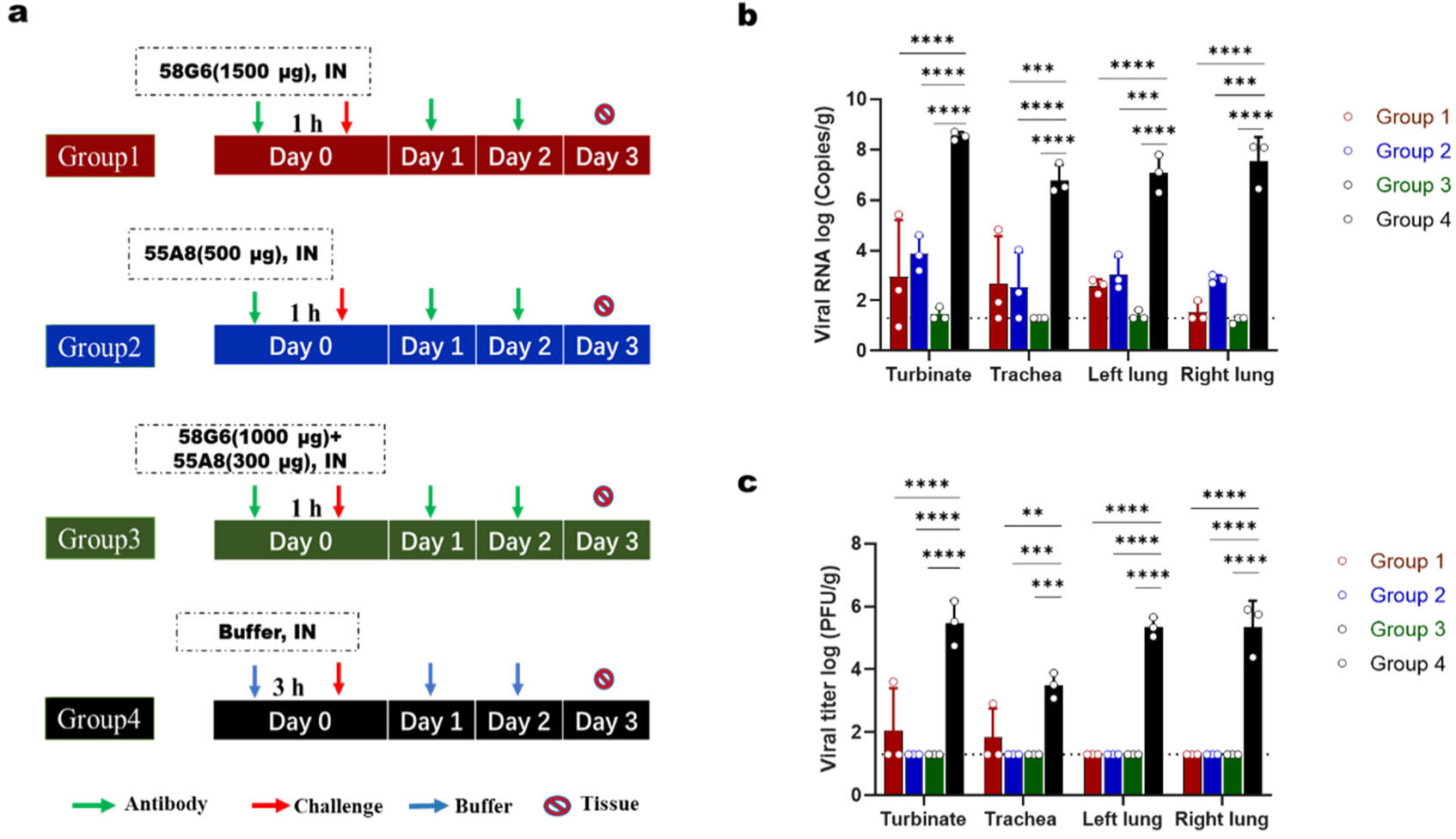
Protective efficacy of 55A8, 58G6 and 55A8/58G6 cocktails against Omicron in the hamster model. Syrian golden hamsters challenged with 10^4^ PFU of Omicron were treated with 58G6, 55A8 or the two-antibody cocktail at 1 h pre-infection and 24 and 48 h post-infection. The turbinates, trachea and lungs were harvested on day 3 post-treatment and analyzed by viral titering (by PFU/g and qRT–PCR). **a** Animal experimental scheme. **b** Viral RNA (log10(RNA copies per g)) was detected in the respiratory tract of hamsters challenged with the Omicron variant at 3 dpi. **c** The number of infectious viruses (PFU) in the respiratory tract was measured at 3 dpi with a viral plaque assay performed with Vero E6 cells.

**Extended Data Figure 4.**
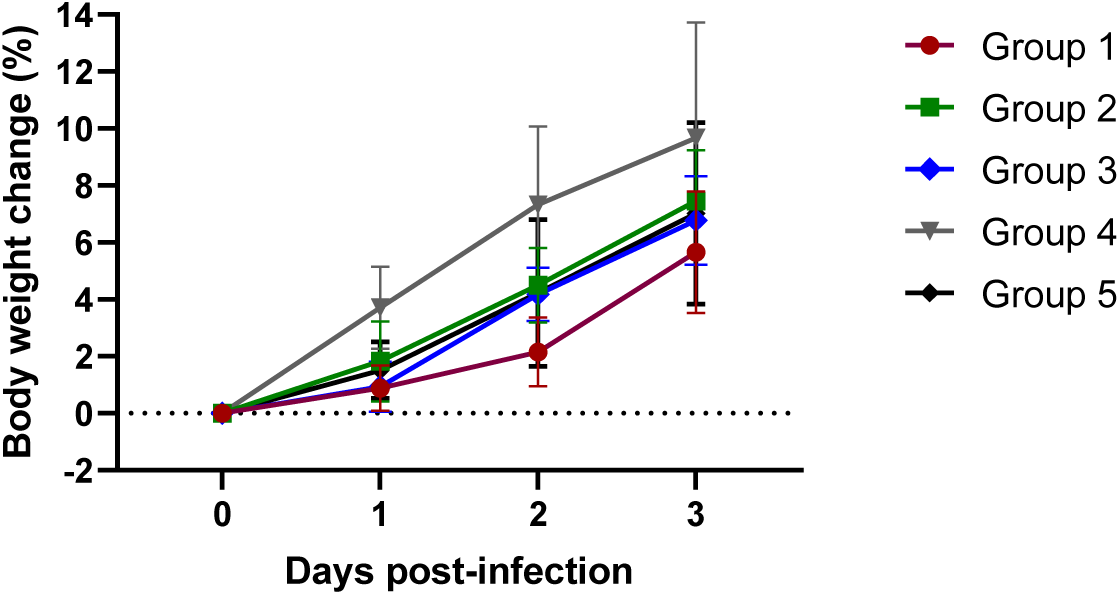
Weight changes in the hamster model treated with 2-cocktail as a measure of the protective efficacy against Omicron.

**Extended Data Figure 5.**
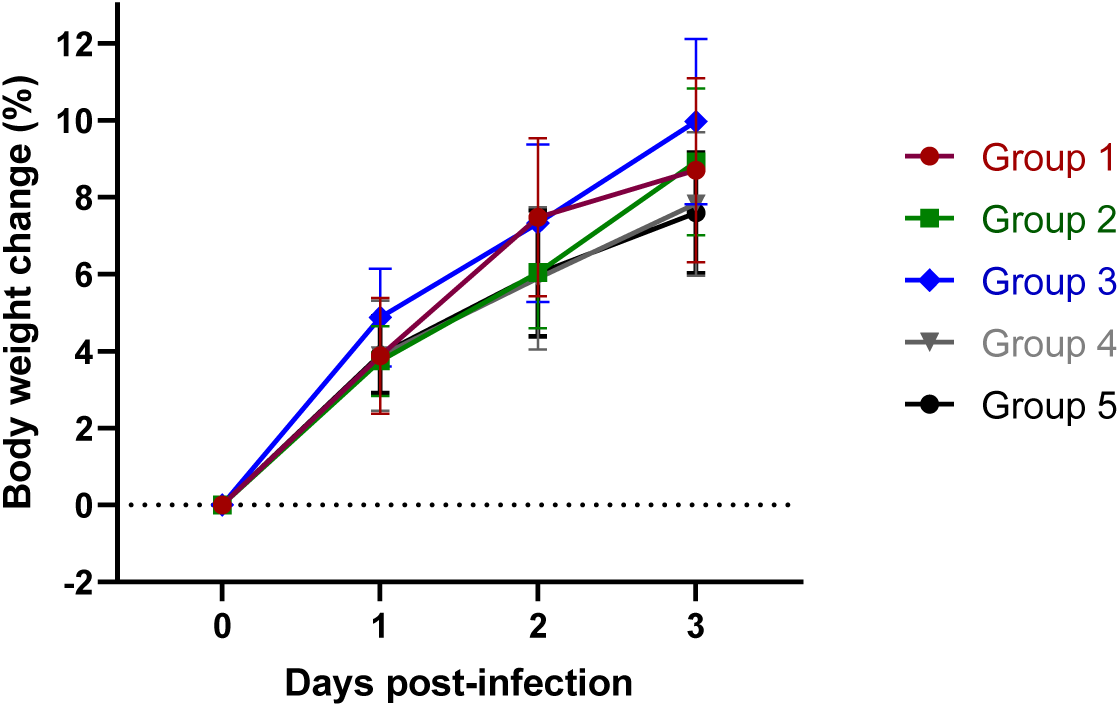
Weight changes in the hamster model after treatment with 2-cocktails as a measure to define the prophylactic dose effective against Omicron.

**Extended Data Figure 6.**
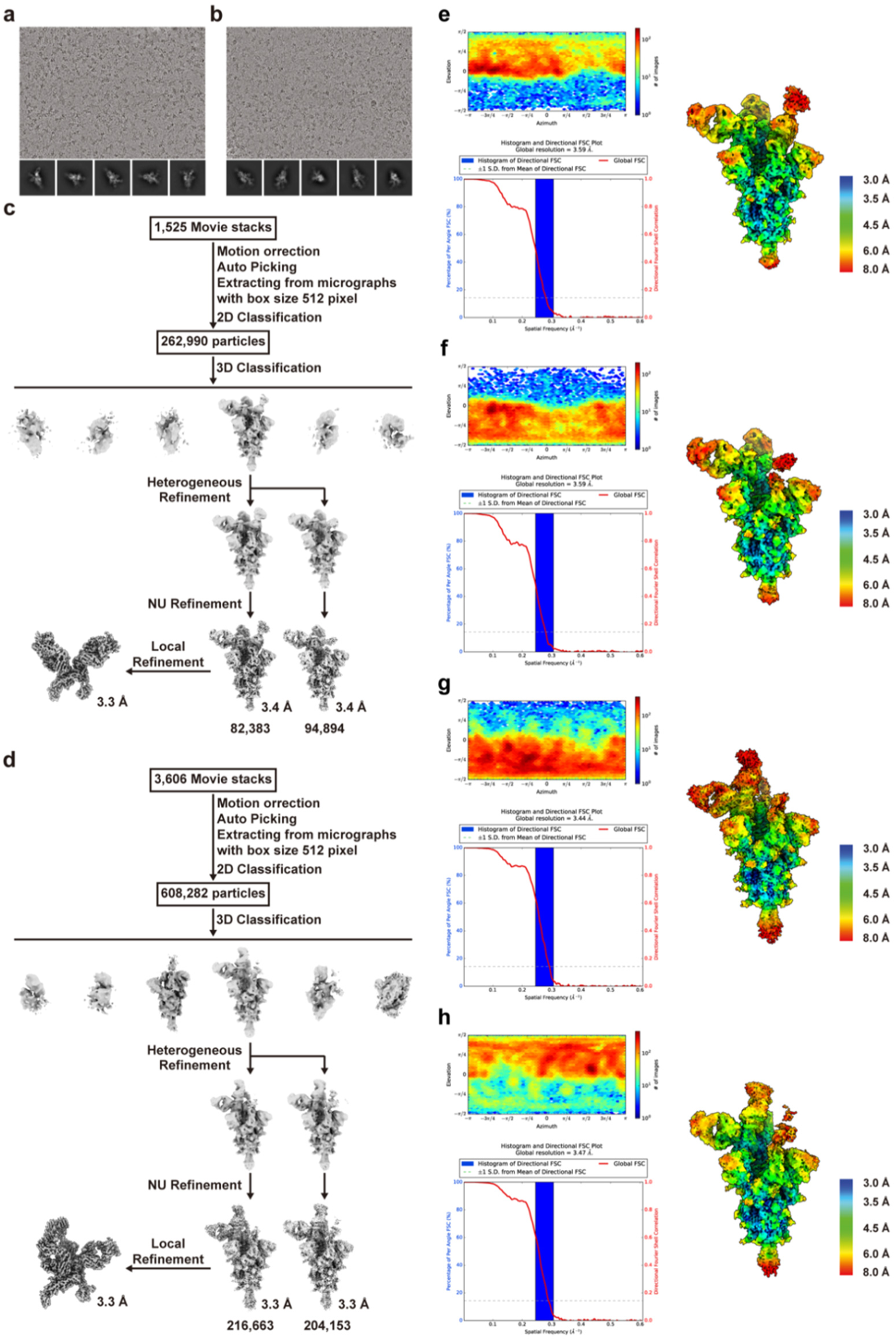
Cryo-EM data processing workflow for Omicron S in complex with the 55A8 Fab or 55A8 Fab-58G6 Fab. a, b Representative cryo-EM micrographs and 2D classes for the Omicron S-55A8 Fab (a) and Omicron S-55A8/58G6 Fab (b) datasets. c, d. Workflow for the Omicron S-55A8 Fab (c) and Omicron S-55A8/58G6 Fab (d) 3D Reconstructions. **e-h** Viewing direction distribution plot, global FSC and histogram, and cryo-EM maps colored by local resolution for the Omicron S-55A8 Fab C ass 1 (**e**) and Class 2 (**f**) and Omicron S-55A8/58G6 Fab Class 3 (**g**) and Class 4 (**h**) datasets.

**Extended Data Figure 7.**
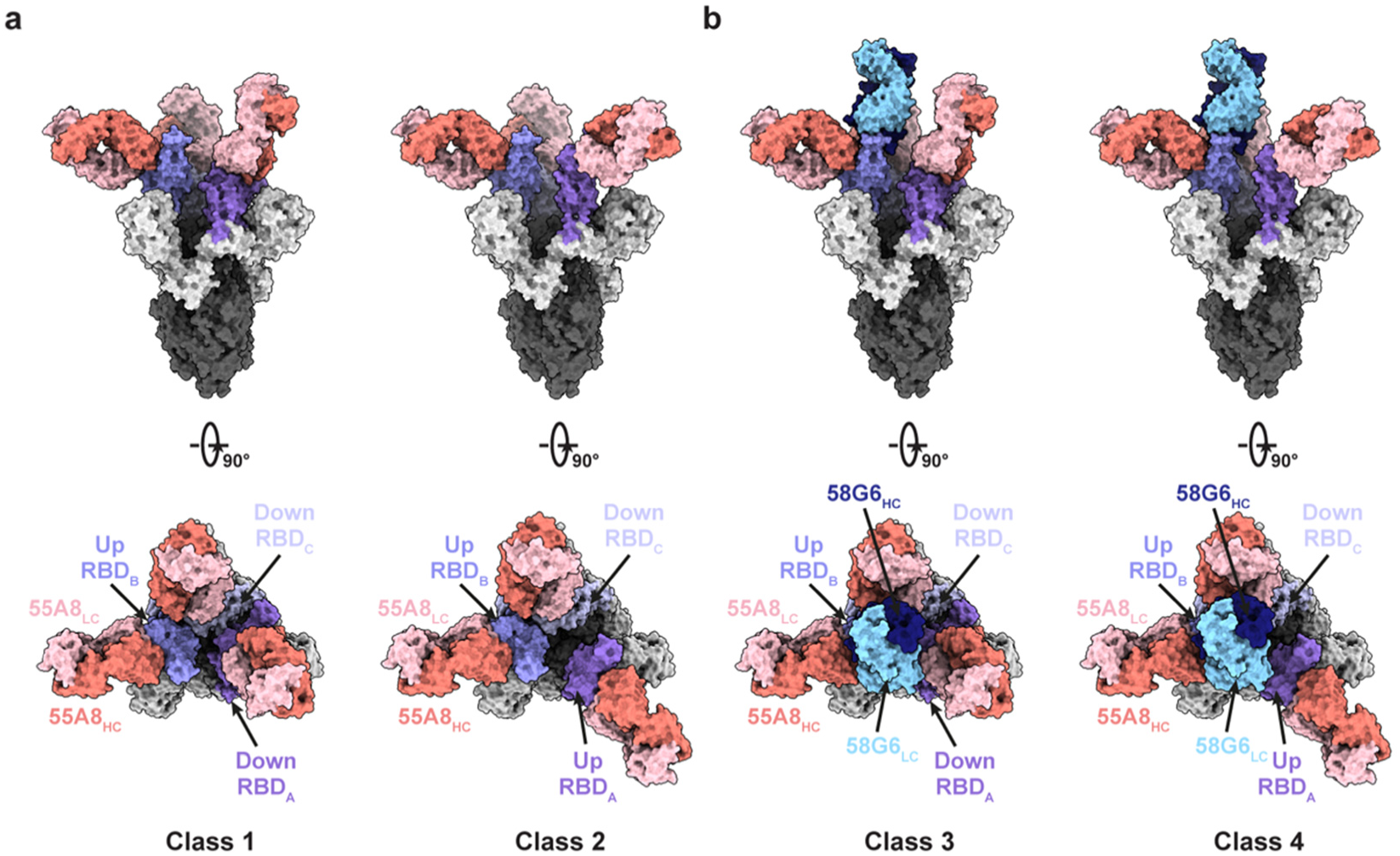
Cryo-EM structures of the Omicron S trimer in complex with 55A8 and 55A8/58G6 cocktails. **a** 55A8 Fabs bind to Omicron S trimers in 2 states. Two perpendicular views of Omicron S-55A8 complexes are shown as the surface. **b** 55A8 and 58G6 Fabs simultaneously bind to Omicron S trimers in 2 states. Two perpendicular views of Omicron S-55A8/58G6 complexes are shown as the surface. 55A8 heavy chain: salmon, 55A8 light chain: pink, 58G6 heavy chain: navy, 58G6 light chain: sky blue, three Omicron RBDs: different shades of purple.

**Extended Data Figure 8.**
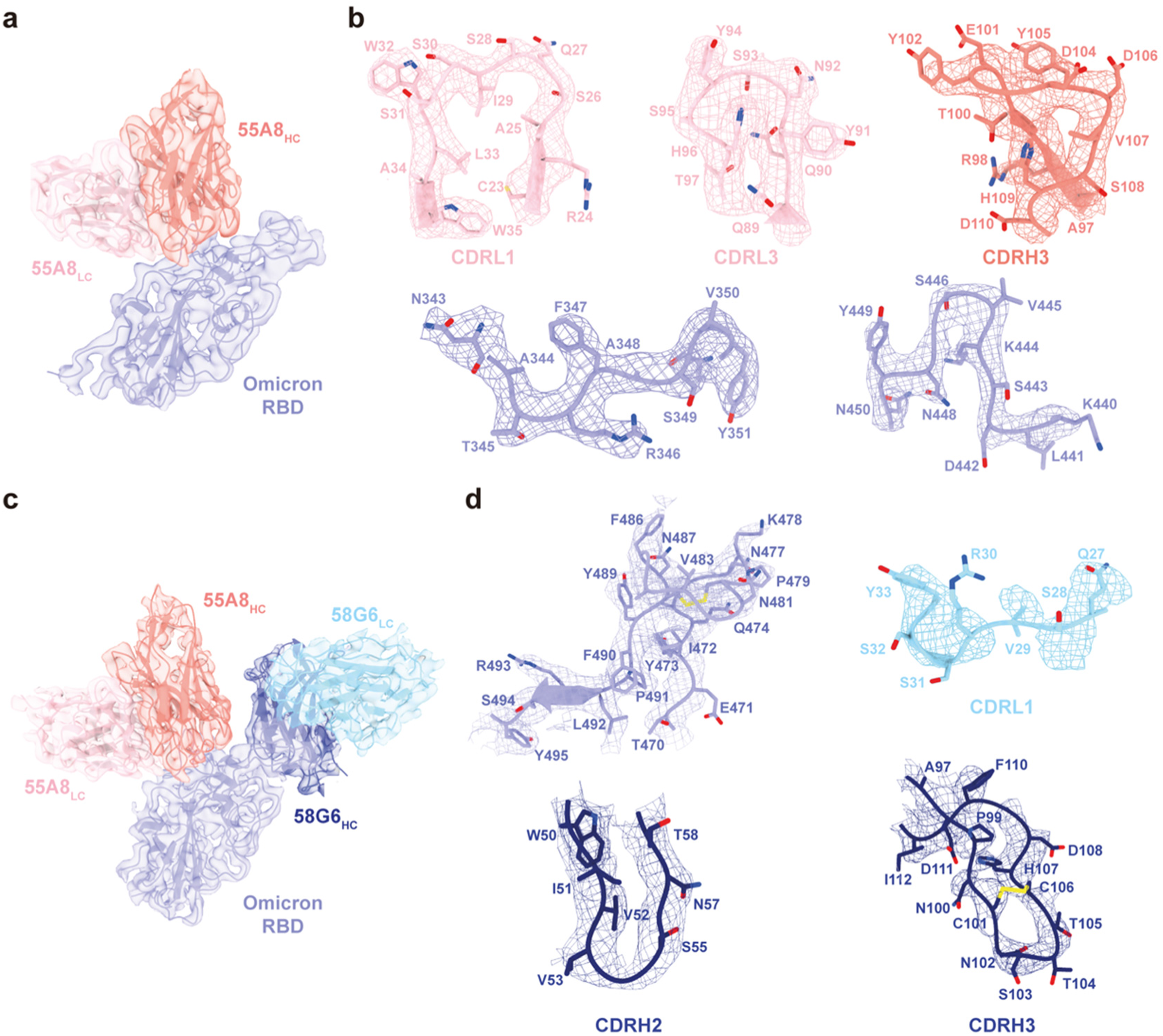
Density maps and atomic models. **a, c** Cryo-EM maps of the binding interface between the Omicron RBD and 55A8 Fab (**a**) or 55A8/58G6 Fab (**c**) variable domains. **b, d** Density maps (mesh) and related 55A8 CDR (**b**) or 58G6 CDR (**d**) atomic models. Residues are shown as sticks, oxygen atoms are colored red, nitrogen atoms are colored blue, and sulfur atoms are colored yellow.

**Extended Data Table 1.** Cryo-EM data collection, model refinement and validation statistics for the Omicron S-55A8 Fab datasets.

|  | 1-up/2-down Class 1 | 2-up/1-down Class 2 | 55A8-RBD Local |
| --- | --- | --- | --- |
| **Data collection and processing** |  |  |  |
| **Voltage (kV)** | 300 | 300 | 300 |
| **Detector** | K3 | K3 | K3 |
| **Pixel size (Å)** | 0.82 | 0.82 | 0.82 |
| **Electron dose (e<sup>-</sup>/Å<sup>2</sup>)** | 60 | 60 | 60 |
| **Defocus range** | -1.2 to -2.8 | -1.2 to -2.8 | -1.2 to -2.8 |
| **Final particles** | 82,383 | 94,894 | 181,072 |
| **Final resolution (Å)** | 3.41 | 3.39 | 3.36 |
| **Model refinement** |  |  |  |
| **Map-model CC (mask)** | 0.86 | 0.87 | 0.87 |
| **Initial model used** | 7WS5 | 7WS5 | 7WS6 |
| **RMSD** |  |  |  |
| Bond lengths (Å) | 0.003 | 0.003 | 0.004 |
| Bond angles (°) | 0.571 | 0.561 | 0.628 |
| Molprobability score | 1.58 | 1.65 | 1.66 |
| Clash score | 7.60 | 8.47 | 6.84 |
| Rotamer outliers (%) | 0.00 | 0.00 | 0.00 |
| Cβ outliers (%) | 0.00 | 0.00 | 0.00 |
| CaBLAM outliers (%) | 2.41 | 2.66 | 2.68 |
| **Ramachandran statistics** |  |  |  |
| Favored (%) | 97.07 | 96.82 | 95.92 |
| Allowed (%) | 2.93 | 3.18 | 4.08 |
| Outliers (%) | 0.00 | 0.00 | 0.00 |

**Extended Data Table 2.** Cryo-EM data collection, model refinement and validation statistics for the Omicron S-55A8 Fab-58G6 Fab datasets.

|  | 1-up/2-down Class 3 | 2-up/1-down Class 4 | 55A8/58G6-RBD Local |
| --- | --- | --- | --- |
| **Data collection and processing** |  |  |  |
| **Voltage (kV)** | 300 | 300 | 300 |
| **Detector** | K3 | K3 | K3 |
| **Pixel size (Å)** | 0.82 | 0.82 | 0.82 |
| **Electron dose (e<sup>-</sup>/Å<sup>2</sup>)** | 60 | 60 | 60 |
| **Defocus range** | -1.2 to -2.8 | -1.2 to -2.8 | -1.2 to -2.8 |
| **Final particles** | 216,663 | 204,153 | 354,108 |
| **Final resolution (Å)** | 3.29 | 3.30 | 3.27 |
| **Model refinement** |  |  |  |
| **Map-model CC (mask)** | 0.84 | 0.88 | 0.82 |
| **Initial model used** | 7WS5 | 7WS5 | 7WS6 |
| **RMSD** |  |  |  |
| Bond lengths (Å) | 0.003 | 0.004 | 0.003 |
| Bond angles (°) | 0.581 | 0.551 | 0.563 |
| Molprobability score | 1.64 | 1.56 | 1.78 |
| Clash score | 8.72 | 6.79 | 7.60 |
| Rotamer outliers (%) | 0.00 | 0.00 | 0.00 |
| Cβ outliers (%) | 0.00 | 0.00 | 0.00 |
| CaBLAM outliers (%) | 2.40 | 2.40 | 2.69 |
| **Ramachandran statistics** |  |  |  |
| Favored (%) | 96.99 | 96.91 | 94.71 |
| Allowed (%) | 3.01 | 3.09 | 5.29 |
| Outliers (%) | 0.00 | 0.00 | 0.00 |

